# On Post-Acquisition Motion Compensation for Prostate Perfusion Analysis

**DOI:** 10.1101/037986

**Authors:** Gert Wollny, Isabel Casanova, María-J. Ledesma-Carbayo, Andrés Santos

**Author notes:** This work was executed at the Group of Biomedical Imaging Technologies, Department of Electronic Engineering, ETSIT, Universidad Politecnica de Madrid, Spain, and with the support of Ciber BBN, Spain. Contact.

## Abstract

Dynamic Contrast enhance magnetic resonance imaging has been established as an accurate method to detect and localize prostate cancer. Time series of three-dimensional datasets of the prostate are acquired and used to obtain per-voxel signal-intensity vs. time curves. These are then used to differentiate cancerous from non-cancerous tissue. However, rectal peristalsis and patient movement may result in spatial-mismatching of the serial datasets and therefore, incorrect enhancement curves. In this work, we discuss and test four methods based on image registration to compensate for these movements. These methods include a serial approach that uses the registration of consecutive images and the accumulation of the obtained transformations, an all-to one registration approach, an approach that first aligns a sub-set of images that are already closely aligned, and then uses synthetic references to register the remaining images, and an approach that uses *independent component analysis* (ICA) to create synthetic references and register the images to these. We conclude that the method based on ICA does not provide a viable approach for motion compensation in prostate perfusion imaging, and that the serial approach fails when motion artifacts are present in the series. The other two approaches provide qualitatively pleasing results.

## I. Preliminary Remark

This work is a partial replication of Casanova [1] that has been done in 2011. Note that on one hand, the redistribution of the data used is restricted because of privacy laws, and on the other hand, that the authors no longer have access to this data. Also note that the implementation of the ICA-based motion compensation method [2] used in this work has since been changed in [3] to follow [4]. Nevertheless, since the claims made in the paper are only of a qualitative nature, it should still be possible to replicate the results.

## II. Introduction

Prostate cancer (PCa) is currently the most frequently diagnosed non-cutaneous cancer and the second most important reason for cancer mortality in men. With an early diagnosis a cure can often be achieved. Dynamic Contrast enhance magnetic resonance imaging (DCE-MRI) has been established as an accurate method in the detection and localization of PCa. DCE-MRI is performed by obtaining repeated, fast T1- weighted images before and up to a few minutes after intravenous injection of a gadolinium containing contrast agent. Three-dimensional datasets of the prostate are acquired every few seconds that allows obtaining a per-voxel signal-intensity vs. time curve. Based on this image series, different pharmacokinetic parameters are modeled and used to differentiate between cancerous and non-cancerous tissue.

However, because of the rectal peristalsis and because patients may move during acquisition, movement may be present in the image series resulting in incorrect enhancement curves and pharmacokinetic parameters. Hence, in order to achieve a proper diagnosis, these movements must be corrected for before making a diagnosis. In this work we present and discuss approaches that compensate for this movement and enable a more reliable and automatic analysis perfusion data.

### A. State of the art

Only a few methods have been published that focuse on motion compensation of prostate perfusion images series. Breathing motion compensation in myocardial perfusion imaging, on the other hand, is a widely discussed topic. In both cases, motion compensation phases a similar challenge: A local movement in an image series needs to be compensated for and the images exhibit a local intensity change over time stemming from the perfusion of a contrast agent. However, certain differences between the two types of perfusion also have to be considered: In myocardial perfusion, where the acquisitions times are lower, usually only a movement of the heart and the surrounding tissue is present, but not a movement of the whole body as it can be seen in prostate perfusion series. On the other hand, the intensity changes induced by the perfusion process poses a larger challenge in myocardial perfusion, since the change is a lot stronger when the contrast agent passes through the left and right heart ventricle. In prostate perfusion images, only the contrast enhancement of the prostate needs to be taken into account.

All methods to compensate motion in image series that we will disscuss are based on image registration. Image registration aims at transforming a study image *S* with respect to a reference image *R*, so that structures at the same coordinates in both images finally represent the same object. Most modern registration approaches use measures that are directly based on the voxel intensities to describe correspondence between the structures in both images. Considering that the rectal peristalsis movement is only a local phenomena, any motion compensation algorithm should allow local, and hence, nonlinear transformations as compared to linear and hence global transformations. If only rigid transformations are allowed, then a region of interest around the prostate must be extracted in order to allow the proper compensation of movement that occurs only locally.

One of the main challenges in voxel based image registration is the establishment of a proper mapping of voxel intensities to tissues that allows to map corresponding anatomical structures in the images correctly onto each other. Since in perfusion imaging these intensities change locally over time, the following general approaches for motion compensation may be considered: Either one registers images from different perfusion phases onto each other and takes the intensity change into account by using a voxel similarity measure that accounts for differences in the tissue-intensity mapping, or one establishes a method to create motion-free synthetic image that exhibit a similar tissue-intensity mapping as its corresponding original image and can, therefore, serve as references in image registration. In such a setting the difficulties of dealing with varying tissue-intensity mappings can be avoided.

In the first approach, image from different time points of the perfusion series, are be registered directly. For example, L udemann et al. [5] used *normalized mutual information* and Breeuwer et al. [6] relied on *cross correlation* (CC) to achive motion compensation in perfusion image series. However, these statistical measures still expect a globally consistent material-intensity mapping but the intensity change due to perfusion happens locally, and because of the cancer perfusion may happen at different rates in different areas of the prostate the statistics may not model the intensity-tissue relation properly. To reduce the impact of the intensity change one may also register only images of the series that are in direct temporal succession and then accumulate the obtained transformations to achieve full motion compensation with respect to a reference frame [7], [8]. However, with this method small registration errors may accumulate and if one registration fails, e.g. because an erroneous frame was recorded due to the patient movement, then it is impossible to achieve a motion compensation for the full series.

The alternative to registering images from different perfusion phases – the creation of synthetic references – was, for example, presented by Milles et al. [9]. They used a three component *independent component analysis* to obtain feature images and a mixing matrix of a myocardial perfusion series. By recombining these images, motion-free synthetic references were obtained that exhibit similar intensities like the original images making it possible to optimize the *sum of squared differences* (SSD) cost measure to achieve registration. While their approach only made use of linear registration its extension to non-linear registration is straightforward.

Another approach that uses synthetic references was presented by Li et al. [10]. In their work they use prior knowledge to estimate a *pseudo ground truth* (PGT) as a reference for non-linear registration to reduce motion. Running this two-step scheme of PGT estimation and image registration in a multi-pass fashion eventually lead to full motion compensation. Amongst other properties the PGT estimation assumes that in the optimal case of motion compensation the time-intensity curves are quite smooth and hence their second order derivatives low. However, because of the noise present in the images the creation this smoothness is not well reflected when the time-intensity curve is evaluated pixel wise. Therefore, the generation of a PGT by also smoothing over time may overcompensate resulting in reference images that do not present the level of detail required for proper motion compensation. The method was later extended to guide registration by a segmentation of the left heart ventricle [11] making the algorithm less generic, i.e. not directly applicable to motion compensation in prostate perfusion date.

The approach of Wollny et al. [12] can be seen as a hybrid to the use of complex registration criteria and the creation of synthetic references. First, they selected a subset of images distributed over the whole series that had a high similarity pointing to images that are already well aligned, and these images were then registered to one global reference optimizing a image similarity criterion based on *normalized gradient fields* (NGF). Then, linear interpolation was used to create synthetic references corresponding to their time index and the corresponding images were registered to these images by optimizing the SSD. This method was also applied to prostate perfusion images [13].

Finally, methods for motion compensation in myocardial perfusion have been proposed that require a segmentation of the left heart ventricle. These methods usually rely on prior knowledge of the heart anatomy that obviously can not be applied to prostate images and are, therefore, not considered here.

### B. Our contribution

In this paper we will analyze the applicability of a subset of the methods described above to motion compensation in prostate perfusion imaging. We present results of motion compensation achieved by running serial registrations, all-to-one registrations, the use of pre-aligned subsets, and the use of ICA. In this work, we only put the focus on an qualitative analysis ofthe registration analysis based on visual inspection of time-profiles and videos. In addition, we provide a free and open source implementation of all the tested methods [3].

## III. Methods

### A. Non-linear registration

Given a *d*-dimensional domain Ω ⊂ ℝ^*d*^ and a space of images 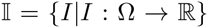, and given a study image *S* ∈ 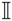 and a reference image *R* ∈ 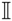, registration aims at transforming the study image *S* with respect to the reference image *R*, so that structures at the same coordinates in both images represent the same object. In practice, this is achieved by finding a transformation *T*_reg_ ∈ 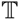 that minimizes a given cost function *F*_cost_: 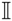 × 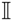 → ℝ, while constraining the transformation through the joint minimization of an energy term *E*: 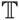 → ℝ:

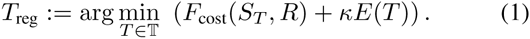

The cost function *F*_cost_ accounts for the mapping of similar structures. *E(T)* ensures topology preservation, which is necessary to maintain structural integrity in the study image, and it thus introduces a smoothness constraint on the transformation *T*. The parameter *κ* is a weighting factor that balances registration accuracy and transformation smoothness.

In nonrigid registration, the transformation *T* needs only to be neighborhood preserving, a restriction that is enforced by the selection of a proper term *E.* In our application, the cost function *F* is derived from a so-called voxel-similarity measure that takes into account the intensities of the whole image domain. As a consequence, the driving force of the registration is calculated directly from the given image data.

Specifically, we employ two image similarity measures: SSD

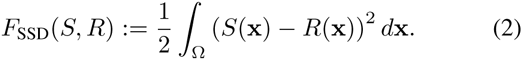

and the formulation of NGF that was given in [12],

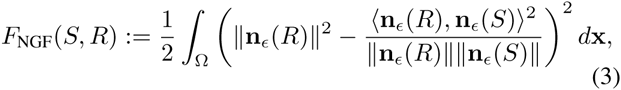

with *d* the number of image dimensions

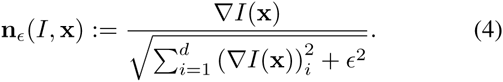

SSD is used when study and reference image exhibit similar intensity distributions – i.e. when synthetic references are used, and NGF is used otherwise. Here, NGF has an advantage over statistical measures like MI or CC, since it is a truly local similarity measure that can accommodate local intensity change. In addition it is fairly easy to implement and as a low computational cost.

The transformation space 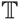 we use is restricted to transformations that can be described by a B-splines based model [14], and we base the regularization on the separate norms of the second derivative of each of the deformation components [15]. The balance between smoothness of the resulting transformation and the amount of non-rigidly that allows for the registration of smaller features can be fine tuned by selecting the B-spline coefficient rate *c*_rate_ and the weighting factor κ. For both parameters, higher values result in smoother transformations that preserve the per-voxel volume better but come at the cost of a reduced ability to register small features.

### B. Motion compensation schemes

The following motion compensation schemes were considered and implemented by using the a freely available toolkit for gray scale image processing [3]:

a) ***Serial***:: Here, only images in temporal succession are registered and then the transformations are applied accumulated. As registration criterion we used a weighted combination of NGF and SSD as proposed in [8].
b) ***AllToOne***:: With this method we register all images to one global reference by using NGF as registration criterion.
c) ***PreAlign***:: The algorithm implements the method proposed in [12] where first a pre-aligned subset of images is estimated. Then, these images are registered non-linearly to a reference picked out of the same subset. Finally, by linearly combining these pre-aligned images synthetic references are created and the remaining images are registered to their synthetic counterparts by optimizing SSD.
d) ***ICARefs***:: Here we took Milles et al. [9] ICA based approach as a blue print and extended by non-linear registration and an alternative method to select the number of independent components as described in [2]. Other then the above algorithms, this algorithm is run in a multi-pass scheme.

## IV. Experiments

### A. Material and Parameters

**Figure 1.**
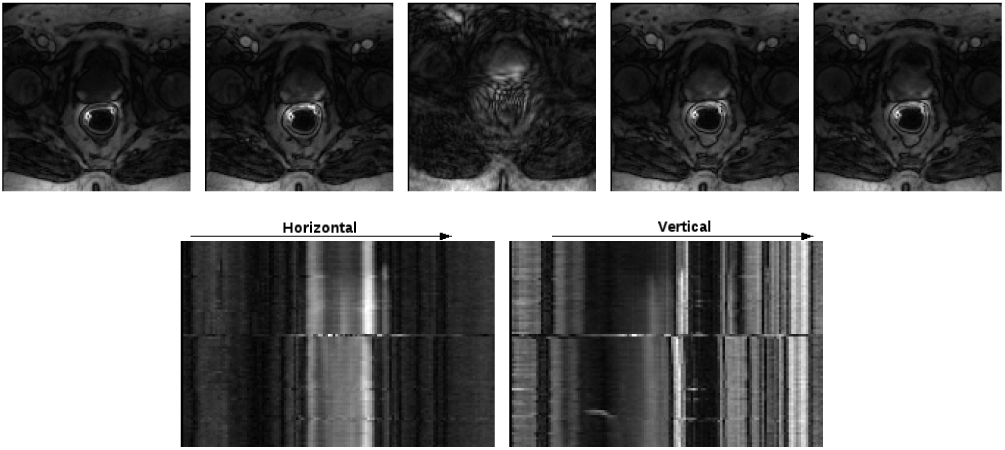
Upper row: example frames of slice ten of patient B. Lower row: the corresponding profiles at the locations indicated in Fig 2, left. Note the motion artifact in the third image which occurs approximately in the middle of the acquisition of the perfusion series as can also be seen in the time-profiles.

Two data sets were acquired from two patients with biopsy proved prostate cancer. For image acquisition a 3T MRI was used in conjunction with a pelvic phased-array coil and an endorectal coil. In order to suppress peristalsis, 40 mg of butylscopolamine was administered intramuscular directly before the MRI scan. The 3D T1-weighted spoiled gradient echo images were acquired directly before and during an intravenous bolus injection of 15 ml paramagnetic gadolinium chelate with an injection rate of 2.5 ml/second followed by a 15 ml saline flush. The series have a temporal resolution of 3 seconds. In one case 110 frames exhibiting perfusion were acquired (Fig. 2), in the other case 84 frames (Fig. 2). The spacial resolution of the images were 1.8mm x 1.8mm x 3mm the images had a size of 128 x 128 x 18 voxels each.

**Figure 2.**
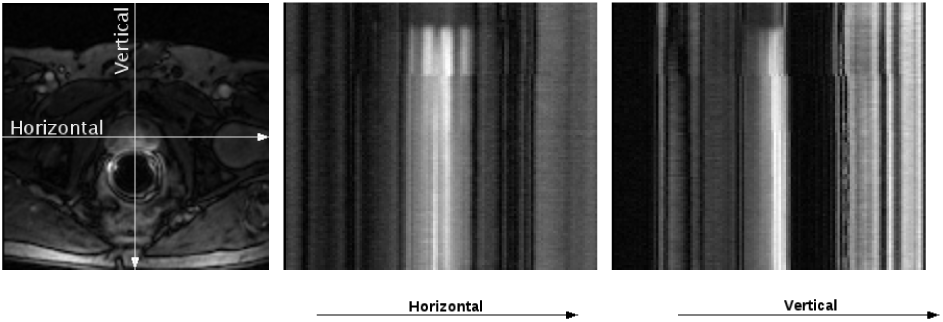
Example frame of the tenth slice ten of patient A that indicates the location of the horizontal and vertical profiles used to assess the quality of the motion compensation (left) and corresponding time-profiles (middle and right).

On both series the motion compensation algorithms described above were executed and the obtained results were qualitatively rated by visually observing videos as well as horizontal and vertical profiles through the time-series stack (see Fig 2, left). The spacial location of these profiles corresponds to row and column number 64 of the tenth axial slice of the data sets.

For the algorithms *Serial*, *AllToOne*, and *PreAlign* the same registration parameters *κ* = 5 and *c*_rate_ = 16 voxels were used, and registration was run by using three multi-resolution levels. Since *ICARefs* is a multi-pass algorithm we used 2 passed and we selected *κ* =10 and *c*_rate_ = 32 in the first pass and reduced these values by the factor of 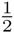 in the second pass. For *Serial* a combined weighted registration criterions of SSD (0.5) and NGF (1.0) was used. In all other case the respective registration criterions were weighted by a factor of 1.0. *PreAlign* has an automatic method for selecting the all-over reference frame. To make comparison of the results easier this frame was then also used as reference with *Serial* and *AllToOne. ICARefs* neither requires the input nor evaluates a specific reference frame.

In all cases we used the rank-1 method of a shifted limited-memory variable-metric algorithm [16] for optimization and the breaking condition were set to an x-tolerance of 0.001, a relative tolerance of the criterion of 0.001, and a maximum of 300 iteration. Optimisation stopped when one of these criteria was fulfilled. All registrations were executed on a AMD Phenom II X6 1035T processor (2.6GHz), Gentoo/Linux 64 bit. The software was compiled with GNU g++ (Gentoo 4.5.2), and optimization flags *-O2-march=native-mtune=native - funroll-loops -ftree-vectorize.*

## V. Results

The per-frame run-time of the used motion compensation schemes was *Serial*=14.5s, *AllToOne*=13.1s, *PreAlign*=17.8s, and *ICARefs*=12.0s. With minor restrictions for *PreAlign* where first a subset of the images has to registered, all registrations can be run in parallel resulting in a near linear speedup when multiple CPUs are utilized.

The methods *Serial*, *AllToOne*, and *PreAlign* all provided comparable good results for patient A (Fig. 3).

**Figure 3.**
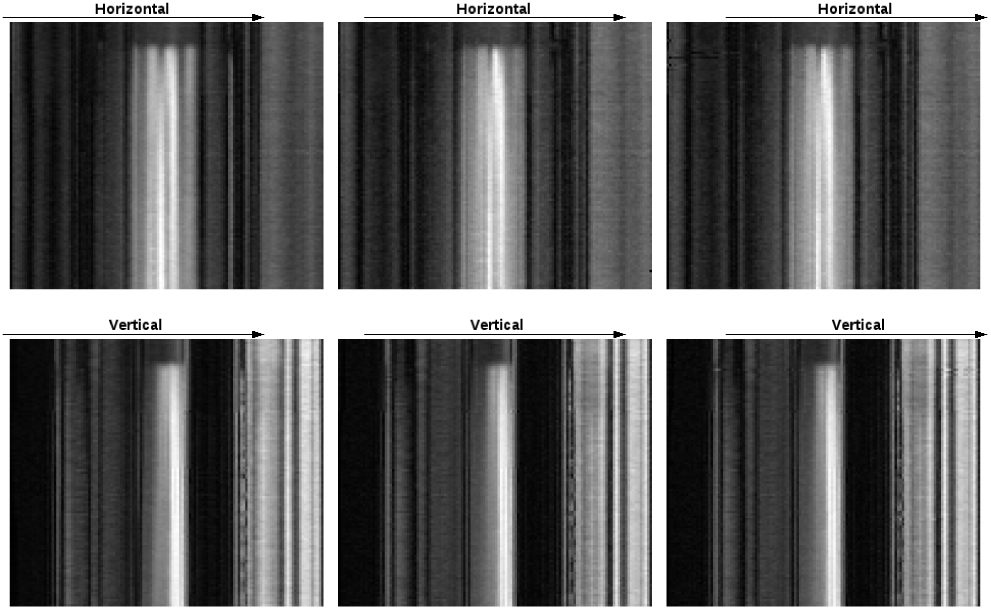
Time-intensity profiles after motion compensation Patient A, horizontal (upper row) and vertical (lower row) profiles of slice ten. From left to right *Serial*, *AllToOne*, and *PreAlign* all provide good motion compensation for this patient.

Since the series of patient B one frame contained a strong motion artifact (Fig. 1, upper row, middle) it was not possible to achieve motion compensation by using *Serial*. With the reference frame located before the frame containing motion and because of the accumulation of the transformations used by this method the information in all frames located after the frame containing the motion artifact are clobbered (Fig. 4).

*AllToOne*, and *PreAlign* again resulted in comparable registration results (Fig. 5), with *PreAlign* reducing the motion at the image boundaries better. This difference between the algorithms can be attributed to the use of different registration criterions. While *PreAlign* used NGF only for some key frames, *PreAlign* relies on this criterion for all image registrations. However, with NGF only gradients contribute to the all-over cost value and, therefore, more local minima may be present in the cost function. With SSD on the other hand, all pixels contribute to the cost value resulting in less local minima and better registration results.

**Figure 4.**
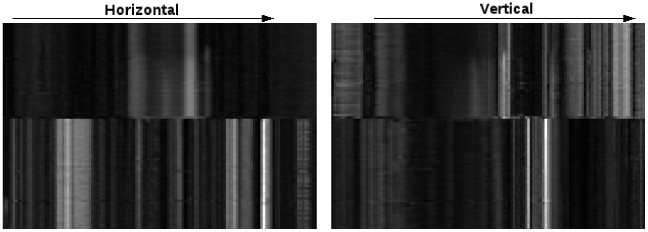
Time-intensity profiles after motion compensation using *Serial*, Patient B. The reference frame is located before the frame exhibiting strong motion and it is clearly visible that no motion compensation could be achieved for the frames after this frame.

**Figure 5.**
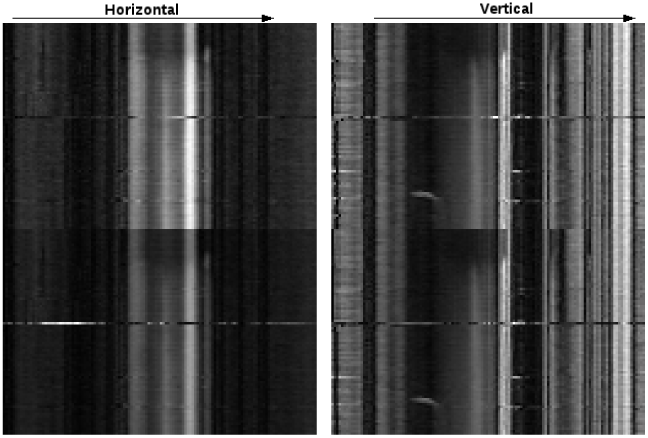
Time-intensity profiles after motion compensation *AllToOne* (upper row), and *PreAlign* (lower row). Generally, a good registration is achieved for both methods, but applying *AllToOne* results in more residual movement close to the image boundaries.

**Figure 6.**
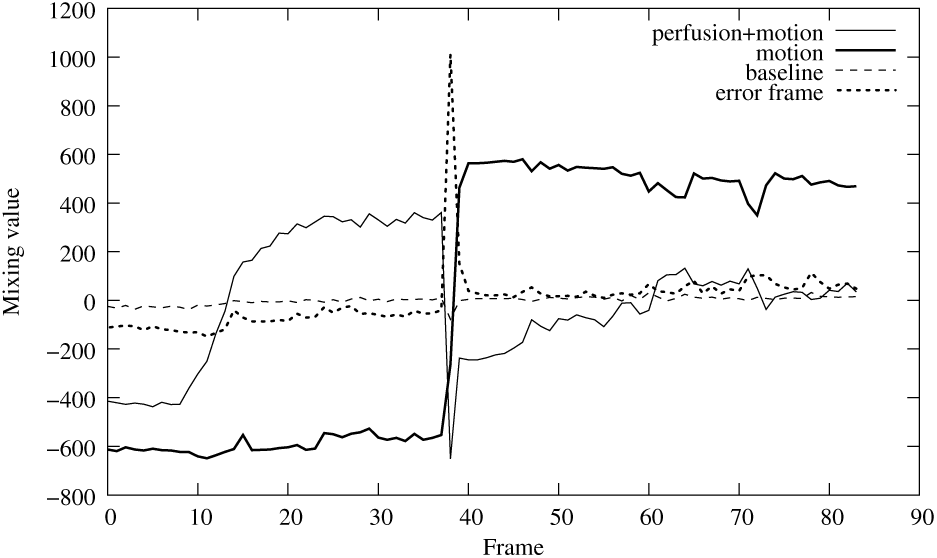
Mixing curves obtained for a four component ICA for Patent B. Note the jumps in the curves *motion* and *perfusion+motion* that relate to the motion component that is fully retained by the ICA. Also note the spike in the curve labeled *error frame* that shows how a four component ICA singles out the frame with the motion artifact as seen in Fig. 1.

With *ICARefs* it was not possible to achieve motion compensation for neither of both data sets. This is because the *independent component analysis*(ICA) retains the motion (see Fig. 6), which is also the reason for the low registration time per frame: since the synthetic references that are very similar to the original images registration is achieved very fast. From Fig. 6 it is also evident that the motion is not separated into one single *independent component*(IC) which makes it impossible to eliminate the motion in the reference images by just removing the corresponding IC from the reference image creation. Note however, that ICA makes it possible to easily identify the frames where strong patient movement actually happens and it also provides information about the intensity change induced by the perfusion. Both types of information could prove helpful when devising new algorithms for motion compensation.

## VI. Conclusion

Out of the tested methods *Serial* motion compensation can be achieved, but only if the images of the perfusion series are free of artifacts, and *ICARefs* motion patterns that relocate the whole imaged body are can not be compensated for.

*AllToOne* and *PreAlign*, on the other hand, proved to be usable for motion compensation of prostate perfusion image series, even when some frames are acquired with strong motion artifacts. Because *PreAlign* needs to evaluate the prealigned subset, *AllToOne* requires less computational time.

Judging based on the visual inspection of the results *PreAlign* provides better results then *AllToOne* – most likely, because for most registrations *PreAlign* uses SSD a registration criterion that has less local minimal then the NGF based measure that is always used with *AllToOne*. In addition, *PreAlign* can be run fully automated, with *AllToOne* one has at least to check that the indicated global reference frame is not actually a frame that was recorded with strong artifacts.

When using *PreAlign* for motion compensation in myocardial perfusion, Wollny et al. [12] reported difficulties in the creation of synthetic references because the linear interpolation used to create synthetic references was not always able to model the strong intensity changes induced by the contrast agent passing through the left and right heart ventricle. In prostate imaging the intensity changes observed are not as intense, and hence the creation of synthetic references does not suffer from this problem.

Future work will include the quantitative confirmation of above results by running a thorough validation based on the comparison of manually obtained time – intensity curves with automatically obtained ones. Finally, one may consider the use of information that can be drawn from an ICA to improve above algorithms.

## VII. Acknowledgment

The authors would like to thank Thomas Hambrock from the Department of Radiology, Radboud University Nijmegen Medical Centre, Nijmegen, The Netherlands for providing the data and medical background.

